# Accuracy of functional gene community detection in *Saccharomyces cerevisiae* by maximizing Generalized Modularity Density

**DOI:** 10.1101/2022.12.28.522153

**Authors:** Pramesh Singh, Jiahao Guo, Jing Li, Urminder Singh, Eve Syrkin Wurtele, Kevin E. Bassler

## Abstract

Identifying functionally-cohesive gene communities from large data sets of expression data for individual genes is a key approach to understanding the molecular components of biological processes. Here, we compare the accuracy of twelve different approaches to infer gene co-expression networks and then find gene communities within the networks. Among the approaches used are ones involving a recently developed clustering method that identifies communities by maximizing *Generalized Modularity Density* (*Q_g_*). RNA-Seq data from 691 samples of *S. cerevisiae* (yeast) are analyzed. These data have been obtained from organisms grown under diverse environmental and developmental conditions and encompass varied mutant lines. To assess the accuracy of different approaches, we introduce a statistical measure, the Average Adjusted Rand Index (AARI) score, which compares their results to Gene Ontology (GO) term associations. Inferring gene networks using the *Context Likelihood of Relatedness* (CLR) and subsequently clustering by maximizing Generalized Modularity Density is found to identify the most significant functional communities. Also, to quantify the extent to which the identified communities are biologically relevant, a GO term enrichment analysis is performed. The results indicate that many of the communities found by maximizing Generalized Modularity Density are enriched in genes with known biological functions. Furthermore, some of the communities contain genes of unknown function, enabling inference of potentially novel functional interactions involving these genes. Furthermore, some genes are species-specific orphan genes; assignment of these orphan genes to communities enriched in a particular biological process provides a method to infer the biological process in which they are involved. We focus on a few communities that are highly significantly enriched in a particular biological process, and develop experimentally-testable predictions about the orphan genes in these communities.

**Author summary:** Finding gene communities that are of biological relevance from expression profiles of individual genes is a critical approach to understanding biological processes and their molecular components. Various computational methods have been developed to infer underlying metabolic and regulatory networks and to identify functional communities of genes. Which network inference and clustering methods works best to achieve this goal has largely remained an open question. Here, using genome-wide transcriptomic data for *S. cerevisiae*, we systematically compare the effectiveness of several commonly used network inference and clustering methods. We rank these methods by comparing the clusters obtained by different methods to Gene Ontology (GO) terms. We find that inferring gene networks using a method known as the Context Likelihood of Relatedness (CLR) and subsequently clustering by maximizing Generalized Modularity Density identifies the most significant functional communities.

## Introduction

Despite the recent surge in abundance of genome-wide biological data and computational capacity, functional annotation of genes still remains a challenging task [1, 2, 3]. Gene metabolic and regulatory networks contain information about interactions among particular genes. Gene communities, also referred to as “clusters” or, when derived from very large diverse data, “regulons”, are groups of coexpressed genes. A community of genes that displays similar expression patterns across a very wide range of experimental conditions is likely to contain co-regulated genes and genes participating in the same biological process; thus identifying gene communities is extremely important to understand their functions [5, 6, 7, 4, 8, 9]. Network-based analyses of gene expression have led to hypotheses on the roles of genes of completely unknown function, which were later experimentally-verified [15, 10, 11, 12, 13, 14].

The first step to finding communities of related genes from gene expression data is to infer a network from the data in which the genes are nodes, and the strengths of the links between pairs of genes are imputed based on some measure of the co-expression of genes. A variety of approaches have been used to infer gene networks based on correlations, such as Pearson’s or Spearman’s correlations, of RNA abundance [15, 16, 17, 4]. Pearson’s and Spearman’s Correlations distinguish positive and negative gene interactions, providing important biological information. However, these measures miss non-linear relationships among gene expression. Information-theoretic methods that use Mutual Information (MI) between gene pairs as a similarity measure have been used to capture these non-linear variations, which can help uncover strong pairwise relationships between genes that are not detected by linear measures [18]. Context Likelihood of Relatedness (CLR), an algorithm for unsupervised network inference [19, 20], uses the *distribution* of the MI of two genes combined with the *value* of the MI [18] between these two genes to compute a *Relatedness* [19] score (see Materials and Methods).

Once a network is created from expression data, a network clustering algorithm can be used to find structure within the network by grouping strongly-connected genes together into communities. Several methods for finding optimal clustering have been developed, each with a somewhat different underlying notion about communities [21, 4, 22]. Even in cases where a community is mathematically well-defined, most commonly-used algorithms to find the optimal clustering are stochastic, i.e., each individual realization of that algorithm finds a slightly different network partition. Here we consider four promising clustering methods: Markov Cluster Algorithm (MCL) [22], and three modularity-based approaches [23, 29, 30]. Each of these methods of partitioning has its pros and cons.

MCL [22] is a widely used method for finding communities in a network; it identifies these communities by a steady-state distribution of simulated random flows in the network. MCL is scalable and fast. However, it requires the specification of external parameters (inflation, expansion). A proper choice of these parameters is important for the algorithm to converge. There is no definite way to select these parameters to ensure convergence. Other limitations of MCL include unknown convergence time and its unsuitability for application to networks with large diameters [22], where the diameter of a network is the longest of all shortest paths between pairs of nodes. MCL has provided a powerful approach to infer the biological function of novel genes, which have later been experimentally confirmed [4, 31, 12].

Modularity (*Q*) [23] is a measure used to quantify community structure in a network. For a given network partition into communities, Modularity is defined as the difference between the fraction of links inside communities and the expected fraction if links were distributed randomly according to a null model. The partition that has the maximum Modularity value corresponds to “the” community structure. However, finding the exact maximum Modularity partition is a computationally challenging, NP-complete problem [24]. Thus, for large networks, such as those analyzed in this paper, it is necessary to use computational algorithms that find approximate solutions that complete in a time that scales as a polynomial in network size. Fortunately, fast, “polynomial-time” algorithms have been developed that can reliably find partitions with Modularity values close to the maximum in networks of up to tens of thousands of nodes [25, 26, 27]. Despite its usefulness, detecting communities by maximizing Modularity has limitations. In particular, small communities in some large networks can not be found with the approach. This issue is known as the “resolution limit problem” [28]. Recently, two new measures, Excess Modularity Density (*Q_x_*) [29] and Generalized Modularity Density (*Q_g_*) [30], have been proposed to overcome this issue. Identifying the community structure of a network with partitions that these modularity density measures significantly mitigates the resolution limit problem. Also, the fast algorithms cited above for finding partitions that maximize Modularity can be adapted to maximize these two modularity density measures.

The goal of this paper is to compare the accuracy of commonly used and the newly developed network inference and clustering methods in detecting gene communities that represent specific biological functions. As a case study, we apply the methods to analyze expression data from a model species, *Saccharomyces cerevisiae* (yeast). RNA abundance profiles generated from raw RNA-Seq data representing the expression of 6692 annotated genes across 691 runs from 44 studies are analyzed, including samples representing a wide range of developmental conditions, strains, growth media, and times [32]. These data are used to construct gene co-expression networks using (a) Pearson’s pairwise correlations, (b) mutual information (MI), and (c) CLR approaches. Then, we use four community detection methods to find communities of interest within the different inferred networks: (i) Markov Clustering (MCL), (ii) maximizing Modularity, (iii) maximizing Excess Modularity Density, and (iv) maximizing Generalized Modularity Density. The clusters found by each of these methods are compared to the *S. cerevisiae* Gene Ontology (GO) leaf terms, i.e. GO terms having no children [33]. The statistical overrepresentation of genes in the clusters is determined using the hypergeomteric test. By comparing gene communities identified by each method with the GO term assigned to each gene, we find that communities found by maxi-mizing Generalized Modularity Density are the most accurate, regardless of which network inference method is used.

## Materials and Methods

### RNA-Seq processing pipeline

The R package *SRAmetadb* was used to query the NCBI-SRA (National Center for Biotechnology Information-Sequence Read Archive) database for publicly available *S. cerevisiae* RNA-Seq runs using the filters: taxon ID = 4932, platform = illumina, and layout = paired. Results with library strategy miRNA-seq, ncRNA-seq, or RIP-seq were filtered out, leaving 691 RNA-Seq samples [32] (see Supplemental Material for full metadata). The *SRA toolkit* was used to download the raw RNA-seq data from the NCBI-SRA database. We used kallisto [34] to quantify the expression of all Saccharomyces Genome Database (SGD) annotated genes (6,692; annotation version R64-1-1) downloaded from ftp://ftp.ensembl.org/pub/release-90/fasta/saccharomyces_cerevisiae/cdna/Saccharomyces_cerevisiae.R64-1-1.cdna.all.fa.gz. To correct for library size effects, we used the Trimmed Mean of M values (TMM) normalization from the *edgeR* (R package) [35] to normalize data between samples followed by Transcripts Per Million (TPM) normalization within each sample. The code/scripts are available at https://bioconductor.org/packages/release/bioc/html/edgeR.html.

### Network inference methods

Network inference involves finding the similarity or relation between the expression profiles for each pair of genes and assigning a score that acts as the link weight between that pair of genes. We used three different measures to allocate weights between genes.

(a) Pearson’s correlation *R*(*X*, *Y*) between genes *X* and *Y* was calculated using the expression profiles of *X* and *Y*.

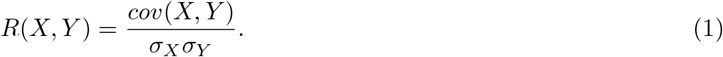

Here *cov*(*X*,*Y*) is the covariance between the expression profiles of gene X and Y, and σ_*X*_ and σ_*Y*_ are their corresponding standard deviations.

(b) Mutual information (MI) between genes X and Y defined as follows

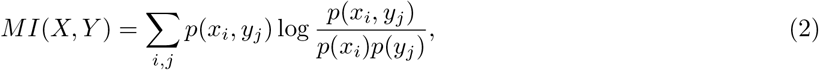

where *p*(*x_i_*,*y_j_*) is the joint probability that gene X has expression level *x_i_* and gene Y has expression level *y_j_*. *p*(*x_i_*) and *p*(*y_j_*) denote the marginal probabilities of gene X having expression level *x_i_* and gene Y having expression level *y_j_* respectively. These probabilities are computed from the original expression profiles using a B-spline smoothing and discretization method. Here, we used the open uniform knot vector with the 10 bins and a spline order of 3 [18]. The mutual information itself is used to define a weighted network where links are weighted by *MI*(*X*,*Y*).

(c) Context Likelihood of Relatedness (CLR) which calculates *relatedness f*(*X*,*Y*) by comparing this mutual information between gene pair X and Y, to the marginal distributions of mutual information for X and Y respectively, which provides a third measure to construct the network. In other words, *MI*(*X*, *Y*) is compared to the distribution of *MI* between gene *X* (or *Y*) and all other genes {*MI* (*X*,*Y*); ∀*Y* (or*X*)}. A Z-score is then calculated using:

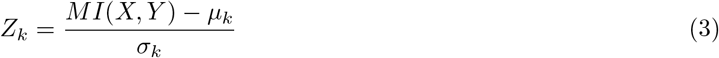

where *μ_k_* and *σ_k_* are the mean and standard deviation of the corresponding distribution. The subscript *k* refers to either gene *X* or gene *Y*. Negative Z-scores are set to be zero. Then the final relatedness value is defined as:

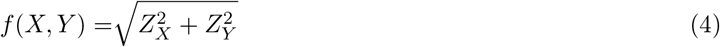

These inferred scores (*R*, *MI*, *f*) can be used as the link weight to connect the corresponding gene pairs. This, however, gives rise to a very dense network with a large number of low-weight links. All three measures can independently be used to infer the gene regulatory network. The observed non-trivial dependencies between these measures for *S. cerevisiae* are plotted in Supplementary Fig. S1.

It is often helpful to discard weaker (low-weight) links by choosing a reasonable threshold for discard; weights for the remaining links can be maintained or set to 1 thus creating a binary network [20]. As a final step, we refined each of the three networks by discarding weaker links from the network. The thresholds for different measures are chosen around a value at which the network begins to fall apart and also keeping in mind that the largest connected component of the network after the threshold is applied is of similar size for the network obtained by different measures (Supplementary Fig. S2).

### Community detection methods

We use four different methods for finding communities in the network. The first method, *Markov Cluster Algorithm* (MCL), computes communities by using steady-state probabilities of a random walk process on the network [22]. Since the communities in a network are characterized by regions of high link density, typically a random walker spends more time within the same community and visits across communities are less frequent. We choose the inflation and expansion parameters as 2. This leads to a steady-state probability distribution which is used to assign communities.

The other three methods we use are based on the detection of modular structures in a network by finding the partition that maximizes a particular measure. Calculating this partition is a computationally challenging task and thus requires efficient computational algorithms [25, 20, 26, 27]. We use the popular *modularity* measure *Q* [23], which is defined as

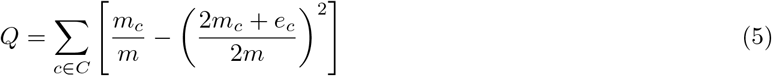

Here *m_c_* is sum of link weights in community *c*, *e_c_* is the weight sum of external links of *c*, and *m* is the total weight of all links in the network.

A notable drawback of modularity is the “resolution limit” (RL) problem [28], which prevents it from detecting small communities in a large network. To enable small clusters to be inferred we have partitioned by maximizing the *Excess Modularity Density* (*Q_x_*) [29] measure, which largely mitigates the RL problem of modularity. The measure, *Q_x_*, for a weighted network can be written as

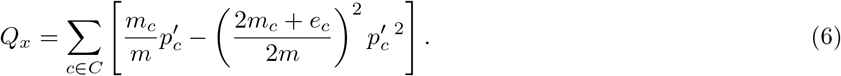

Here *m_c_* is sum of link weights in community *c*, *e_c_* is the weight sum of external links of *c*, and *m* is the total weight of all links in the network. 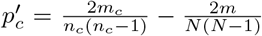 measures the excess density of links inside *c*, where *n_c_* is the number of nodes in *c* and *N* is the network size.

Finally, we partition by maximizinng *Generalized Modularity Density Q_g_* [30], which was recently developed to eliminate the RL problem, allowing detection of communities at any desired resolution. Thus, the measure *Q_g_* is particularly desirable for studying hierarchical community structure in biological networks, and identifying functional gene communities that are hidden in other modularity approaches because of their small size. The Generalized Modularity Density measure is defined as

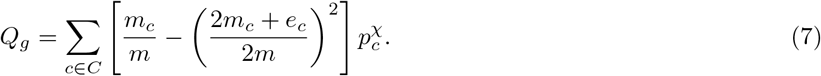

where *χ* is an exponent that controls the granularity of the clusters, and 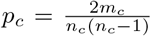 is the link density of community *c*. In the limit *χ* = 0, this metric reduces to modularity *Q* and larger values of *χ* result in more fine-grained clusters. For this research, we choose *χ* =1 such that the link density factor is linear.

We used a variant of the algorithm prescribed in [26] to detect communities by maximizing *Q_x_* [29]. Maximization of *Q* and *Q_g_* was performed by the Reduced Network Extremal Ensemble Learning (RenEEL) scheme [27, 30].

### Average Adjusted Rand Index (AARI)

Several methods exist for comparing network partitions. Adjusted Rand Index (ARI) [36], a variant of Rand Index [37] that has been adjusted for random chance, is a popular method for comparing the similarity of two non-overlapping partitions of a network. In contrast, Omega Index [38] compares two overlapping partitions but is less useful in comparing a partition with overlapping communities with one in which all communities are non-overlapping. Therefore, we selected ARI to compare the relative efficacy of various gene expression network partitions to the “ground truth” GO term associations [33]. In our case, the gene communities that we detect are non-overlapping (each gene has only one community membership). In contrast, the grouping of genes given by GO term associations is an overlapping partition because a gene can have multiple GO term associations. We define a measure to compare such partitions. First, we turn the overlapping GO term partition into a multiple non-overlapping partitions by considering each node belonging to one GO term at a time. For a small example network with overlapping partition, the procedure is visualized in Fig. 1. We then compare the original non-overlapping partition to every extracted non-overlapping GO term partition one by one and compute the ARI. Ideally, we would like to average over all of the individual ARIs to compute an average ARI.

**Figure 1:**
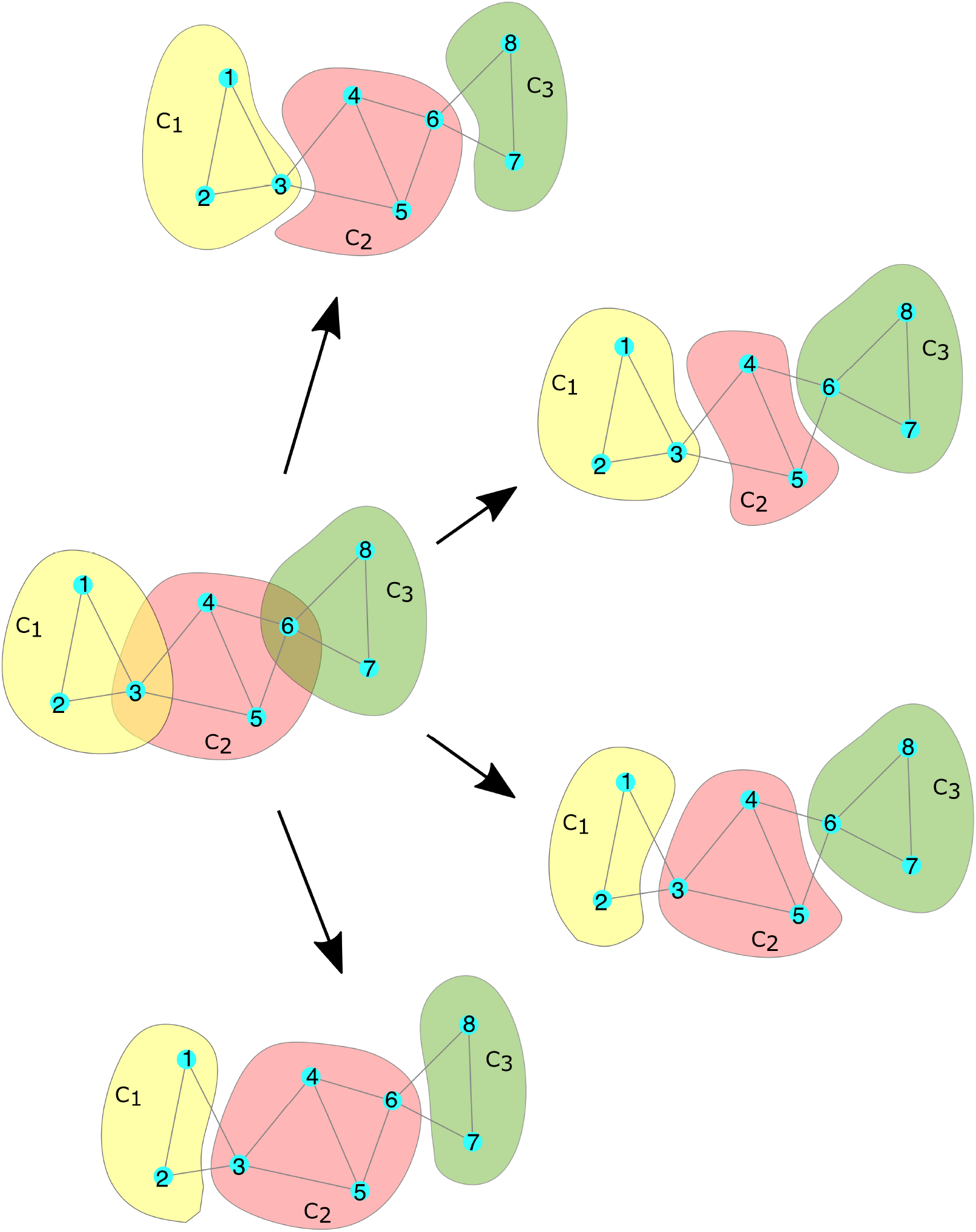
Extracting non-overlapping partitions from an overlapping partition. A simple example of a network consisting of 8 nodes (labeled 1 through 8). The three communities, represented by the colored blobs, are labeled as *C*_1_,*C*_2_,*C*_3_. The initial overlapping partition (center) of this example network, in which node 3 and node 6 each belong to two communities, is used to construct four non-overlapping partitions by allowing each node to be a member of only one community. Each of the four possible extracted non-overlapping partitions (periphery) is indicated by an arrow.

This is computationally infeasible because of the huge number of non-overlapping partitions that can be extracted from a partition even with moderate overlap. The exact number of possible non-overlapping partitions is given by 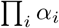, where for every gene *i*, *α_i_* represents the number of GO terms that it is associated with. To meet this challenge, we randomly select a finite but sufficiently large ensemble of *N_S_* non-overlapping GO term partitions from the original overlapping GO term partition. We then use the average ARI over this ensemble to compare the non-overlapping gene communities with GO term associations.

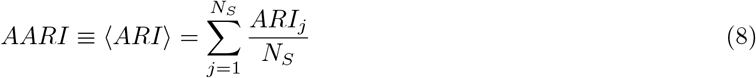

where *N_S_* is the number of partitions in the ensemble and *ARI_j_* is the ARI with respect to a given sample. For a large enough *N_S_*, the distribution of *ARI* converges and according to the *Central Limit Theorem* [39], the *AARI* over the ensemble 〈*ART*〉 provides a good approximation for average overall implicit partitions. A summary of these steps is shown in Fig. 2.

**Figure 2:**
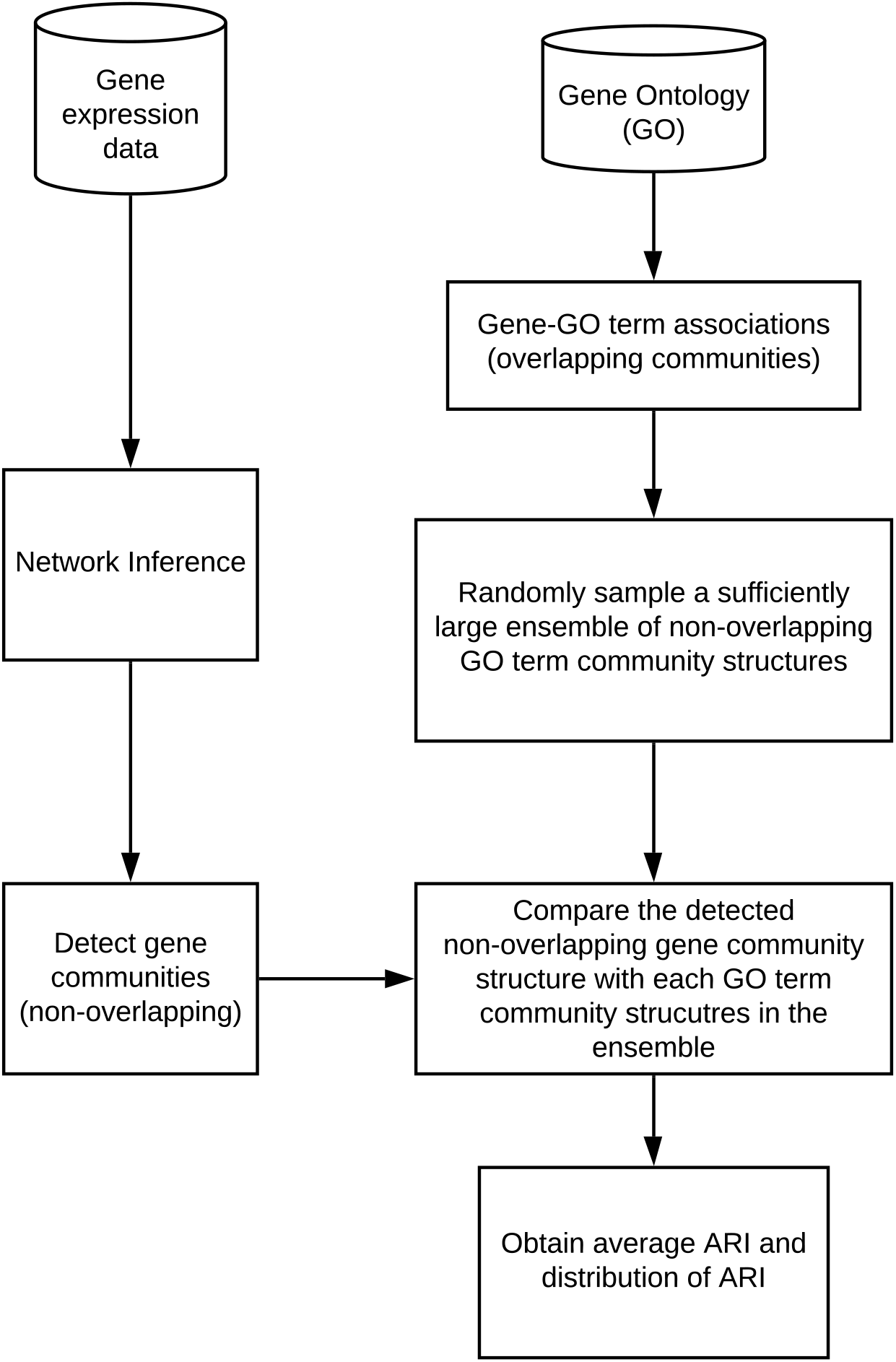
Steps to compute the AARI score. A co-expression network is inferred from the gene expression data by a network inference method. This inferred network is partitioned into non-overlapping communities using a community detection algorithm. Finally, the resultant partition is compared with each partition contained in the ensemble of non-overlapping partitions extracted from the overlapping gene-GO term associations. ARIs are calculated for each partition, and then the average ARI is computed.

### GO term enrichment

For the assessment of Gene Ontology term enrichment in communities, we considered leaf GO terms only [33]. Leaf GO terms are not further subdivided, and thus they correspond to the most specific biological functions. The enrichment analysis uses the hypergeometric test that computes the p-value for overrepresentation. The p-values are adjusted by applying the Benjamini-Hochberg correction [41] for multiple testing. The p-value for the hypergeometric test is given by

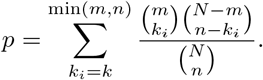

Here, *n* is the number of genes in the community, *m* is the number of genes in the GO term, *k* is the number of genes common between the community and the GO term, and *N* is the total number of genes. We used the open source tool *Ontologizer* [40] for this purpose.

## Results

### Network inference

We use three different inference methods: Pearson correlation (*R*), Mutual Information (*MI*), and Relatedness (*f*), to construct weighted networks from the gene expression data. Each network consists of weighted links given by the specific measure *R*, *MI*, or *f*. Threshold values are chosen to create a similar-sized largest connected network for each inference method: 0.7 for Pearson correlation network, 0.26 for Mutual Information network, and 3.0 for Relatedness network (red dashed lines in Supplementary Fig. S2). The Weights below the threshold are set to 0 and the weights above the thresholds are retained except when applying MCL. For MCL, we construct binary networks, where these weights are set to 1 above the threshold and 0 below the threshold.

### Gene communities

We used four community detection methods to detect community structure in each network. These methods are MCL, and community detection by *Q*, *Q_x_*, and *Q_g_*. Each network inference approach combined with a different community detection method results in a different partition of genes into communities, i.e., four different community detection methods and three different networks, yielding twelve sets of communities.

To determine which combination of network inference method and community detection method is more relevant biologically, we compare our results to the Gene Ontology terms. Specifically, we compare all twelve partitions to the structure of GO terms using the AARI (Fig. 3 (a)). First, we observe that all community detection methods show improved performance when applied to Relatedness network constructed by the CLR method. Other network inference methods showed similar and relatively low performance. Second, we find that the results from the Relatedness network combined with Generalized Modularity Density (denoted by *Q_g_* (*f*)) shows a better match (higher AARI) with the GO terms than other methods (although closely followed by *Q_x_*(*f*)). We also show the overall distribution of ARI when using the Relatedness network in Fig 3(b) to show that *Q_g_* consistently outperforms *Q*, *Q_x_* and MCL. See Supplementary Material for a complete list of genes and their community membership as assigned by *Q_g_* applied to the Relatedness network.

**Figure 3:**
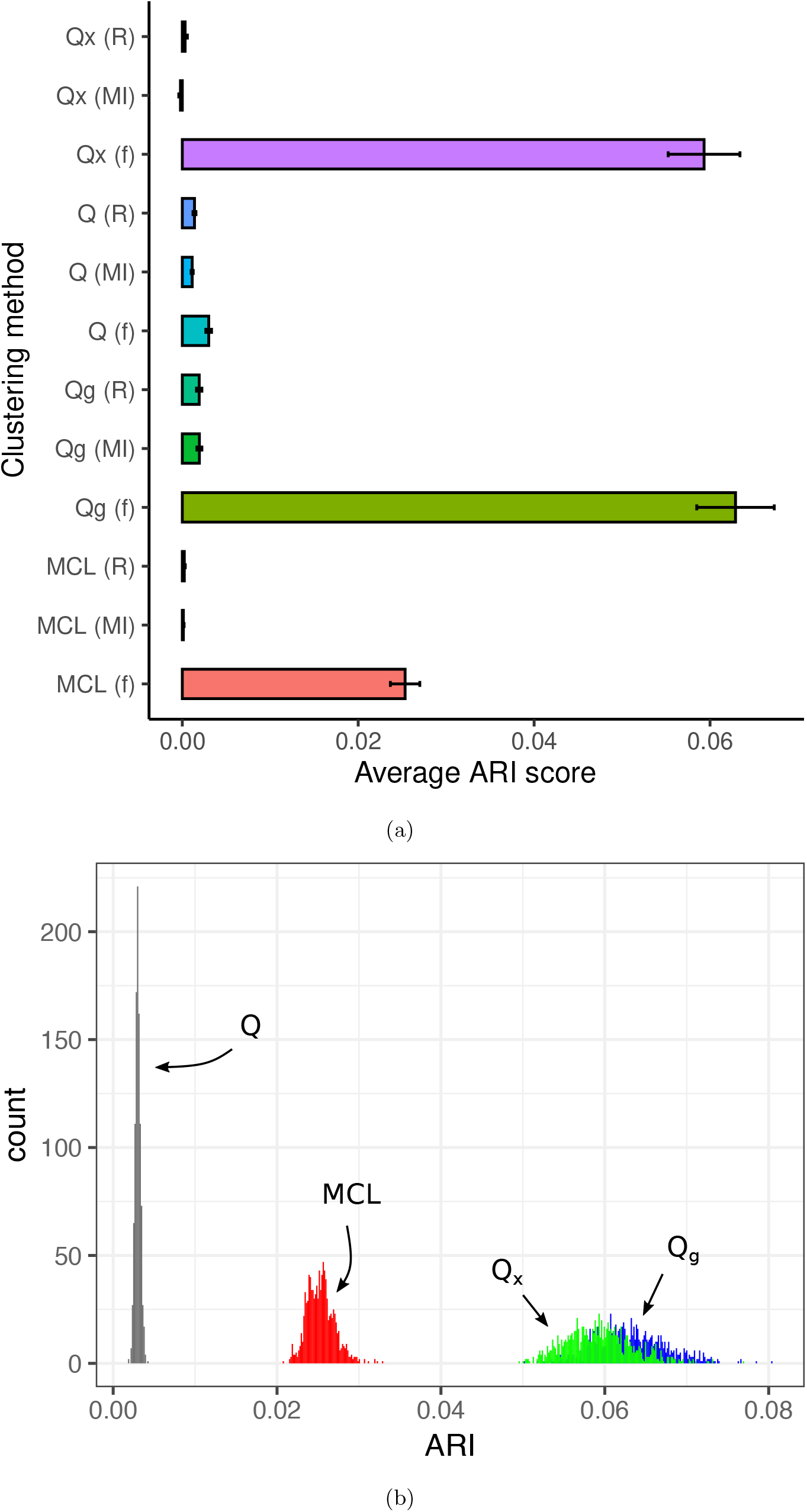
Comparison of GO term associations and gene communities identified by different network inference and community detection methods. (a) Average Adjusted Rand Index (AARI) scores for 1000 realizations comparing GO-terms and communities found by applying modularity *Q*, excess modularity density *Q_x_*, Markov Clustering (MCL), and Generalized Modularity Density *Q_g_* to networks determined by Relatedness (*f*), Mutual Information (*MI*), and Pearson correlation (*R*). A higher ARI indicates a better match. Error bars show one standard deviation interval around the mean. (b) The distribution of ARI scores of gene communities and GO terms over 1000 realizations identified by Modularity *Q*, Excess Modularity density *Q_x_*, MCL, and Generalized Modularity Density *Q_g_*.

### GO-term enrichment

The comparison of four clustering methods on networks inferred using three inference methods revealed that the best community inference was obtained by using Generalized Modularity Density *Q_g_* on the Relatedness network. Further, we focus on communities of *S. cerevisiae* obtained by this method.

Table 1 shows the ten communities with the most significant (smallest p-values) GO-term enrichment. Communities were obtained by applying metric *Q_g_* to the Relatedness network. Table 2 shows the top ten matches of the same enrichment analysis for communities that contain orphan genes. Orphan genes are genes encoding species-specific proteins. They are present across eukaryotics and prokaryotics, and are thought to play an important role in speciation [42, 43, 44, 45, 46]. The functions of the vast majority of orphan genes remain unknown. Of those orphan genes that have been researched, many are implicated in defense and offense/predation, either as secreted molecules, or as interactors of internal defense responses [43, 46, 47, 49].

**Table 1:** GO term enrichment of gene communities containing orphan genes, as identified from RNA-Seq data by creating a gene expression matrix by Context Likelihood of Relatedness and clustered using Generalized Modularity Density. The ten most significant associations are shown.

| community | GO terms | genes in community | genes in GO term | genes in common | p-value | orphans |
| --- | --- | --- | --- | --- | --- | --- |
| 5 | structural constituent of ribosome | 191 | 220 | 129 | 1.10E-152 | 0 |
| 70 | RNA-DNA hybrid ribonuclease activity | 55 | 48 | 40 | 2.66E-80 | 0 |
| 70 | DNA-directed DNA polymerase activity | 55 | 61 | 40 | 4.15E-73 | 0 |
| 70 | ATP binding | 55 | 664 | 42 | 8.45E-29 | 0 |
| 27 | endonucleolytic cleavage in ITS1 to separate SSU-rRNA from 5.8S rRNA and LSU-rRNA from tricistronic rRNA transcript (SSU-rRNA, 5.8S rRNA, LSU-rRNA) | 181 | 43 | 25 | 2.34E-26 | 0 |
| 27 | endonucleolytic cleavage to generate mature 5'-end of SSU-rRNA from (SSU-rRNA, 5.8S rRNA, LSU-rRNA) | 181 | 32 | 22 | 1.56E-25 | 0 |
| 27 | endonucleolytic cleavage in 5'-ETS of tricistronic rRNA transcript (SSU-rRNA, 5.8S rRNA, LSU-rRNA) | 181 | 31 | 21 | 3.74E-24 | 0 |
| 5 | rRNA export from nucleus | 191 | 18 | 15 | 3.05E-19 | 0 |
| 112 | structural constituent of ribosome | 17 | 220 | 15 | 8.06E-19 | 0 |
| 1 | cytochrome-c oxidase activity | 19 | 20 | 8 | 7.28E-15 | 9 |

**Table 2:** GO term enrichment of gene communities containing orphan genes, as identified from RNA-Seq data by creating a gene expression matrix by Context Likelihood of Relatedness and Generalized Modularity Density. Ten most significant associations are shown.

| community | GO term | genes in community | genes in GO term | genes in common | p-value | orphans |
| --- | --- | --- | --- | --- | --- | --- |
| 1 | cytochrome-c oxidase activity | 19 | 20 | 8 | 7.28E-15 | 9 |
| 1 | heme binding | 19 | 27 | 5 | 9.74E-08 | 9 |
| 1 | Group I intron splicing | 19 | 12 | 4 | 2.04E-07 | 9 |
| 1 | mitochondrial electron transport, cytochrome c to oxygen | 19 | 14 | 3 | 2.68E-05 | 9 |
| 34 | carnitine O-acetyltransferase activity | 187 | 3 | 3 | 0.0010 | 5 |
| 1 | proton-transporting ATP synthase activity, rotational mechanism | 19 | 17 | 2 | 0.0027 | 9 |
| 34 | fructose 2,6-bisphosphate metabolic process | 187 | 4 | 3 | 0.0032 | 5 |
| 1 | homing of group II introns | 19 | 1 | 1 | 0.0062 | 9 |
| 1 | movement of group I intron | 19 | 1 | 1 | 0.0062 | 9 |
| 66 | protein targeting to vacuole involved in ubiquitin-dependent protein catabolic process via the multivesicular body sorting pathway | 15 | 22 | 3 | 0.0074 | 6 |

Orphans provide a disruptive force in evolution and may be expressed sparsely and in distinct patterns from those of other genes [46, 43]. However, identifying orphans that are community members provides an avenue towards building inferences about their function. The three communities CLR detected that contained multiple orphan genes are: *Cluster 1*, structure and assembly of the cytochrome C oxidase subunit of mitochondrial aerobic respiration, and intron splicing of cytochrome C oxidase mitochondrially-encoded genes [50]; *Cluster 34*, intermediate metabolism; *Cluster 66*, multivesicular body sorting associated with ubiquitin-mediated protein catabolism. A detailed enrichment analysis is included in the Supplemental Material.

## Discussion

A variety of methods for network inference and clustering have been applied to gene expression data. Most have their own advantages and drawbacks, depending on the specific problem at hand and the data type. A previous study using a CLR method demonstrated its effectiveness in inferring known gene interactions [19]; the CLR algorithm performed better than several other approaches in inferring regulatory interactions among genes in microarray expression data from the prokaryote, *E. coli*. Thus, we used this approach in inferring the co-expression network of *S. cerevisiae* genes, along with Pearson correlation and Mutual Information. To our knowledge, it had not yet been applied to a eukaryotic organism, which have a considerably more complex regulatory and metabolic network properties [51]. It also had not been applied to RNA-Seq data. The key advantage of RNA-Seq data is that it enables insight into the accumulation of all transcripts, whereas many transcripts are not represented on microarray chips [52]. Thus, even the levels and patterns of accumulation of unannotated transcripts can be monitored and associated with particular biological processes. However, RNA-Seq determinations are influenced by the sequencing depth, the method of library preparation, and other technical artifacts [53].

Here, we compare commonly-used methods for inferring functional gene communities from RNA-Seq expression data [32] from the model eukaryote, *S. cerevisiae*. First, using the expression profiles of *S. cerevisiae* genes, we obtained three different weighted co-expression networks, each by employing a distinct method for assigning link-weights between pairs for genes. Accurate inference of the co-expression network is an important step since the downstream analyses depend on the structure of this network. Among the three methods (See Materials and Methods), CLR [19] is shown to perform better at inferring regulatory interactions than others. In *E. coli*, the CLR methods is shown to enhance the average precision by 36% relative to the next-best performing algorithm [19]. In our study, we also observe it to be a better network inference algorithm.

We then applied four different community detection algorithms to identify functional modules in each of the inferred co-expression networks. Three of these algorithms are based on maximizing modularity (*Q*) and two modularity density measures (*Q_x_* and *Q_g_*) that were recently introduced [29, 30] to mitigate the resolution limit problem of modularity [28]. MCL is a widely used random-walk based community detection algorithm, which we also included for comparison. A careful evaluation of these clustering methods revealed that generalized modularity density *Q_g_* had the best performance while modularity *Q* performed worse than the other three algorithms. This is because modularity discovers large clusters that do not correlate with specific GO-terms, which, in contrast, are relatively small. While MCL showed performance gain over *Q*, the two modularity density methods significantly higher performance with the Generalized Modularity Density (*Q_g_*) outperforming all other methods. This can be attributed to the fact that *Q_g_* assigns relatively higher weights to denser communities. Although *Q_x_*, which also introduced weights that are functions of internal link density of a community, as pointed out in [30], it shows resolution problems in very sparse networks. On the other hand, *Q_g_* is resolution-limit free and robust to variability in network structures.

Further, the performance of each method is assessed by evaluating the similarity between GO term associations and the detected gene communities, using the Average Adjusted Rand Index (AARI). The GO term associations of genes overlap (even the leaf GO-terms) and the methods considered in this paper detect nonoverlapping communities. Thus, using ARI score (or other similarity measures of the type) could be deceptive. Thus, to fairly assess the communities, we introduced an average ARI score that is computed over an ensemble of standard ARI scores obtained by randomly sampling non-overlapping GO associations (see Materials and methods). While we use the average of ARI scores to assess the similarity of overlapping clusters because it is intuitively easier to interpret, other measures that quantify similarity between two non-overlapping partitions (such as Normalized Mutual Information [54] could also be used, and averaging any such measure over the prescribed ensemble could be carried out in a similar fashion.

## Conclusion

We systematically evaluated twelve different combinations of network construction and clustering approaches. Our investigation empirically shows that the Relatedness network obtained by CLR combined with community detection using the Generalized Modularity Density *Q_g_* metric outperforms other methods. A focused analysis reveals that communities containing orphan genes have highly significant associations with biological processes, as determined by GO term analysis. This promising approach to identifying gene communities can enrich our understanding of the roles of orphan genes and other genes of unknown function, and infer their relationship with other genes.

## Supporting information

Enrichment-Qg(f)

Gene-communities-Qg(f)

## Supporting information

### Files

yeast-SI.pdf

Gene-communities-Qg(f).xlsx

Enrichment-Qg(f).xlsx

### Data and code availability

All data used in this study are publicly available and have been appropriately cited. The codes to construct networks and to perform clustering are available at: https://github.com/prameshsingh/yeast-clustering.

## Acknowledgments

This work was supported by the NSF through grants DMR-1507371 and IOS-1546858. This work used the Extreme Science and Engineering Discovery Environment (XSEDE), which is supported by National Science Foundation grant number ACI-1548562, through allocation TG-MCB190098.

