## Supplemental Figs. S1 and S2 for "Accuracy of functional gene community detection in *Saccharomyces cerevisiae* by maximizing Generalized Modularity Density"

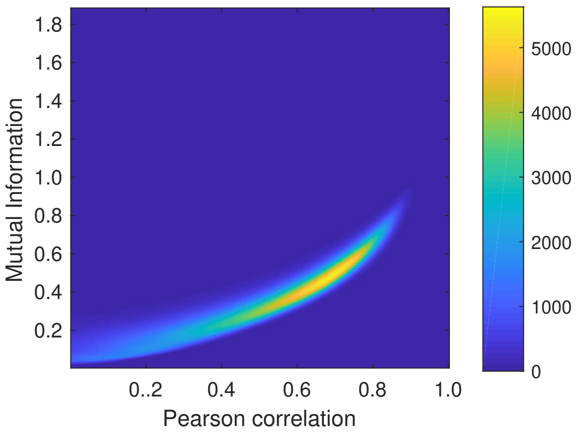


(a)

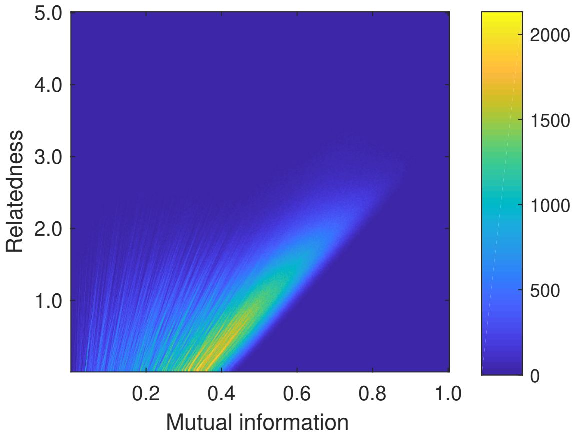


(b)

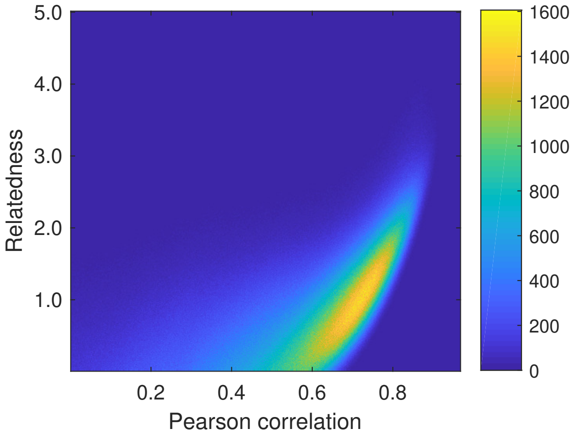


(c)

Figure S1: **Different measures of similarity.** 2-dimensional distribution of link weights for inferred gene co-expression network of *S. cerevisiae*. The area in each figures is divided in to $(500 \times 500)$ bins. The color represents the number of links within each bin. Negative Pearson correlations have been discarded. (a) Mutual information vs. Pearson correlation. (b) Mutual information vs. Relatedness. (c) Relatedness vs. Pearson correlation.

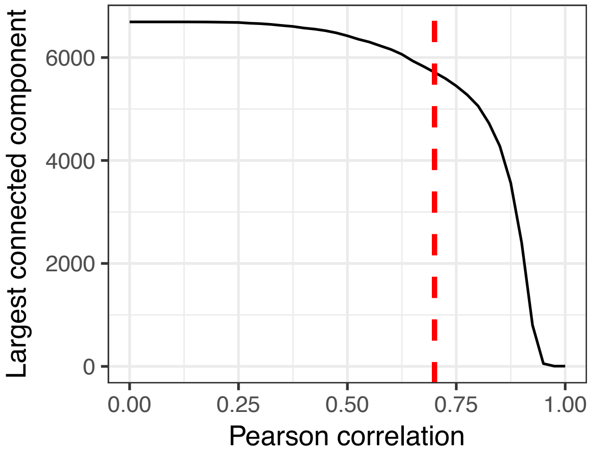


(a)

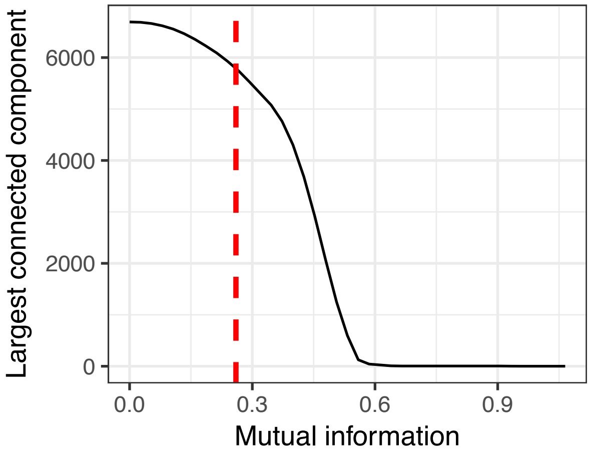


(b)

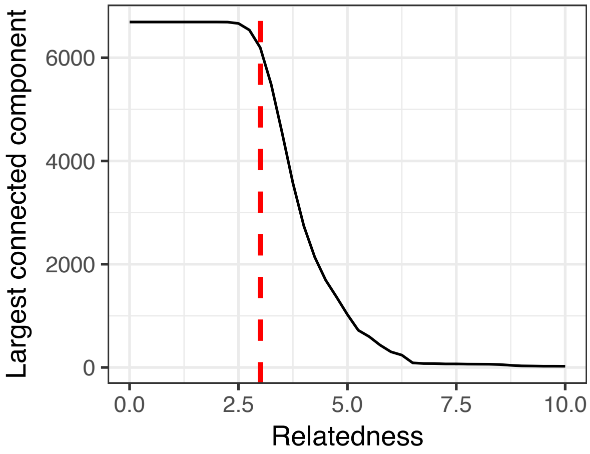


(c)

Figure S2: **Largest connected component of gene networks derived using different methods of network inference.** The largest connected component as a function of threshold for (a) Pearson's correlation network (positive correlations only) (b) mutual information network (c) relatedness network. The threshold for each network (dashed red lines) is chosen such that the size of the largest connected component is approximately same .
